# Evaluation of tumor antigen-specific antibody responses in patients with metastatic triple negative breast cancer treated with cyclophosphamide and pembrolizumab

**DOI:** 10.1101/2022.08.19.504403

**Authors:** Eric D. Routh, Mark G. Woodcock, Jonathan S. Serody, Benjamin G. Vincent

## Abstract

**Background:** The role of B cells in antitumor immunity is becoming increasingly appreciated, as B cell populations have been associated with response to immune checkpoint blockade (ICB) in breast cancer patients and murine models of breast cancer. Deeper understanding of antibody responses to tumor antigens is needed to clarify the function of B cells in determining response to immunotherapy.

**Methods:** We evaluated tumor antigen-specific antibody responses in patients with metastatic triple negative breast cancer treated with pembrolizumab following low dose cyclophosphamide therapy using computational linear epitope prediction and custom peptide microarrays.

**Results:** We found that a minority of predicted linear epitopes were associated with antibody signal, and signal was associated with both neoepitopes and self-peptides. No association was observed between signal presence and subcellular localization or RNA expression of parent proteins. Patient-specific patterns of antibody signal boostability were observed that were independent of clinical response. Intriguingly, measures of cumulative antibody signal intensity relative to immunotherapy treatment showed that the one complete responder in the trial had the greatest increase in total antibody signal, which supports a potential association between ICB-dependent antibody boosting and clinical response. The antibody boost in the complete responder was largely driven by increased levels of IgG specific to a sequence of N-terminal residues in native Epidermal Growth Factor Receptor Pathway Substrate 8 (EPS8) protein, a known oncogene in several cancer types including breast cancer. Structural protein prediction showed that the targeted epitope of EPS8 was in a region of the protein with mixed linear/helical structure, and that this region was solvent-exposed and not predicted to bind to interacting macromolecules.

**Conclusions:** This study highlights the potential importance of the humoral immune response targeting neoepitopes as well as self epitopes in shaping clinical response to immunotherapy.

## BACKGROUND

The successful priming, expansion, and trafficking of neoantigen-specific T cells to the tumor microenvironment is thought to be critical for effective antitumor immunity [1]. T cell killing of tumor cells is a final common pathway of immune surveillance and immunotherapy efficacy [2]. Neoantigen frequency (potentially estimated by tumor mutational burden) may be associated with survival and immunotherapy response [3], as neoantigens can be targets for antitumor T cell responses. Neoantigen-directed therapies including adoptive T cell transfer and therapeutic vaccines are in clinical trials and have been associated with clinical responses in some patients [4-9].

Although T cells have been a central focus of cancer immunotherapy studies, there is a growing appreciation for the role of B cells in anti-tumor immunity. We recently reported that increased B cell gene signature expression and B cell receptor diversity in pretreatment samples were associated with clinical response to immune checkpoint blockade (ICB) in triple negative breast cancer (TNBC) [10]. In murine models of TNBC, ICB response has been found to be dependent upon B cell responses [11]. Beyond TNBC, intratumoral presence of tertiary lymphoid structures (TLS; enriched with B cells, T cells and dendritic cells (DCs)) have been associated with ICB response in various cancer types [12-16]. In ovarian cancer, B-cell-derived IgA production promoted myeloid cell-dependent killing of ovarian cancer cells and antibody-dependent transcriptomic changes in cancer cells that sensitized them to T cell killing [17]. B-cell-derived antibody responses have also been reported to promote the differentiation of neoantigen-specific CD4+ T cells, which can in turn enhance CD8+ T cell effector functionality through IL-21 production [18]. Thus, there is a dynamic interplay between DCs, T cells, and B cells within tumors, and further elucidation of the cellular and molecular mechanisms that govern this dynamic would be valuable for understanding immunotherapy response.

The goal of this study was to understand whether tumor-specific antibodies could be discovered in TNBC patients treated with immunotherapy. To assess this, we used genomics data to predict linear epitopes in eleven patients for generation of custom peptide arrays, which were probed in a multiplex ELISA with patient plasma from two time points: pre-ICB and after two cycles of ICB. We found that a minority of predicted epitopes were associated with IgG antibody signal. Nuclear, cell surface and cytoplasmic locations were predominant in antibody-associated proteins, and RNA expression was not associated with antibody signal. For some patients, including the complete responder, the majority of peptides with antibody signal displayed increased signal after ICB treatment. Furthermore, ICB-dependent boost of both self-peptide-specific and mutated peptide-specific antibody signal was observed for some patients. A set of high-boosted epitopes that were self-specific in the complete responder was observed, and these epitopes were found in an N-terminal, surface-exposed region of the oncogenic EPS8 protein. Together, these data offer an initial glimpse into the characteristics of antibody responses in TNBC patients treated with immunotherapy.

## METHODS

### Whole-exome sequencing and variant calling

WES was performed on FFPE tumor tissue collected prior to treatment on the LCCC1525 trial of low-dose cyclophosphamide plus pembrolizumab in metastatic triple negative breast cancer (NCT02768701), with PBMCs collected serving as the matched normal. Library preparation was performed with the TruSeq DNA, PCR-Free kit (Illumina, San Diego, California, USA) and pooled samples sequenced on the HiSeq4000 platform (Illumina). Somatic and germline WES sequencing files were aligned to Hg38 using bwa (v0.7.17) and sorted, indexed, and duplicates marked using biobambam2 (v2.0.87). BAMs were realigned with Abra2 (V.2.22), followed by somatic and germline variant detection with Strelka2 (V.2.9.10), Cadabra (from Abra2 V.2.22) and Mutect2 (GATK V.4.1.4.0). Capture of exonic sequences was verified using the Picard (V.2.21.1) CollectHsMetrics tool, and quality of sequencing data verified using FastQC (V.0.11.8), and the Picard suite’s CollectAlignmentSummaryMetrics, CollectInsertSizeMetrics, QualityScoreDistribution, and MeanQualityByCycle tools. Variants were filtered by the following criteria: protein-coding mutations only, Cadabra indel quality >10.5, Mutect2 indel quality >6.8 or single nucleotide variant (SNV) quality >9.2, Strelka2 indel quality >15.2 or SNV quality >19.7. Remaining variants required at least five supporting reads and a minimum read depth of 40, or 10 supporting reads and minimum read depth of 80 if MAF <5%. Variants with a MAF >5% in normal tissue were dropped, as were variants appearing at rates above 1% in any subpopulation in either GnomAD or 1000 Genomes databases. To counter FFPE artifacts, C>T and G>A substitutions required a minimum MAF of 10%. Tumor mutational burden (TMB) was calculated from small indels and substitutions identified by WES, and divided by the megabases adequately covered by sequencing reads.

### RNA sequencing

Samples of total RNA extracted from FFPE tumor tissue (ROCHE High Pure FFPE kit, Indianapolis, Indiana, USA) were used to prepare Illumina TruSeq RNA Access (Cat. No. 20020189) sequencing libraries. Sequencing was performed in the UNC-Chapel Hill High Throughput Sequencing Facility on an Illumina HiSeq 4000 platform using the Illumina HiSeq SBS 150 Cycles (PE-410-1001) with 2×75 paired end base reads.

### Neoantigen peptide prediction

HLA major and minor class I alleles were determined from RNA expression data using OptiType v1.3.1 [19] via the authors’ published Docker container in RNA mode (--rna, as per https://github.com/FRED-2/OptiType). Annotated variant transcripts were created using ANNOVAR v2019Oct24 [20], using their suggested SNP filters: the Exome Aggregation Consortium (ExAC) repository (last updated May 16, 2019) and ANNOVAR’s modified dbSNP list, avnsp147. Functional prediction and annotation were done using the suggested dbNSFP v3.0a [21] database. NeoPredPipe v4.0 [22] was used to orchestrate the process of variant filtering, transcript annotation, and binding affinity calculations.

### Peptide selection

Peptide candidates were generated from predicted variant protein sequences (arising from SNVs and INDELs) by iterating over possible 15mers including at least one variant amino acid in a sliding-window approach. For each predicted variant 15mer, a matched reference 15mer was produced as a control. Neoantigen peptides from frameshift or stop-loss mutations were discarded. Single amino acid substitutions were sorted based on their estimated amino acid exchangeability [23], with more dissimilar substitutions favored over more similar ones.

### Peptide arrays

Neoepitopes and matched self-peptides were screened using peptide arrays printed by PEPperPRINT GmbH (Germany). Peptides were converted into two identical microarrays for each patient. The resulting arrays contained varying numbers of linear peptides (range: 2,136-5,498 total peptides, half of which were mutant peptides and the other half comprising matched self-peptides) printed in duplicate, and were framed by additional HA (YPYDVPDYAP, 52 or 40 spots respectively) and polio (KEVPALTAVETGAT, 52 or 38 spots respectively) control peptides. Microarrays were pre-stained with the secondary and control antibodies (Goat anti-human IgG (Fc)-DyLight680 (0.1 µg/ml), Mouse monoclonal anti-HA (12CA5)-DyLight800 (0.1 µg/ml)) in incubation buffer (PBS, pH 7.4 with 0.05% Tween-20 + 10% Rockland blocking buffer MB-070) to investigate background interactions with linear peptides. Subsequent incubation of peptide microarrays with patient plasma samples of the respective patient at 1:20 dilution was followed by staining with secondary and control antibodies. Read-out was performed with an Innopsys InnoScan 710-IR Microarray Scanner at scanning gains of 50/10 (red/green). The additional HA peptides framing the peptide microarrays were simultaneously stained as internal quality control to confirm assay performance and peptide microarray integrity. Quantification of spot intensities and peptide annotation were based on 16-bit gray scale tiff files that exhibit a higher dynamic range than the 24-bit colorized tiff files. Microarray image analysis was done with PepSlide Analyzer. A software algorithm breaks down fluorescence intensities of each spot into raw, foreground and background signal, and calculates averaged median foreground intensities and spot-to-spot deviations of spot duplicates. A maximum spot-to-spot deviation of 40% was tolerated, otherwise the corresponding intensity value was zeroed. For analysis purposes, a background-corrected signal intensity of >500 F.U. (fluorescence units) and less than 2,000 F.U. was denoted as weak signal, and signal intensity >2,000 F.U. was denoted as moderate/strong signal (per recommendations from Pepperprint technical support team).

## RESULTS

### A minority of predicted neoantigen-containing peptides were targeted by endogenous antibodies

We examined IgG reactivity to neoantigen-containing versus unmutated self linear epitopes from patients with TNBC who participated in a clinical trial to examine efficacy of PD-1 inhibition following cyclophosphamide treatment [10] (**Fig. 1**). Of the 40 patients enrolled, eleven patients were chosen for evaluation based on differential response to ICB (1 CR, 4 PR and 6 PD). The predicted peptides were prioritized/ranked (see methods) and were printed on custom peptide arrays along with their corresponding normal self-peptides. Plasma drawn on cycle 1, day 1 (C1D1, pre-study) and cycle 3, day 1 (C3D1, after two completed cycles of pembrolizumab) was used to probe the arrays. Peptide spots that bind antibody would yield fluorescence signal after staining with a secondary antibody conjugated to a fluorescent label. As seen in the digitized scan images of representative microarrays (**Fig. 1**), a large majority of predicted peptides were not associated with IgG antibody binding, and this was true of both mutated and self-peptides. On average, 94.8% of predicted mutant peptides had signal denoted as very weak/noise (<500 F.U.), 3.66% had weak signal (>500 F.U. and <2000 F.U.), 1.24% had moderate signal (>2,000 FU and <10,000 F.U.), and 0.26% had strong signal (>10,000 F.U.).

**Figure 1.**
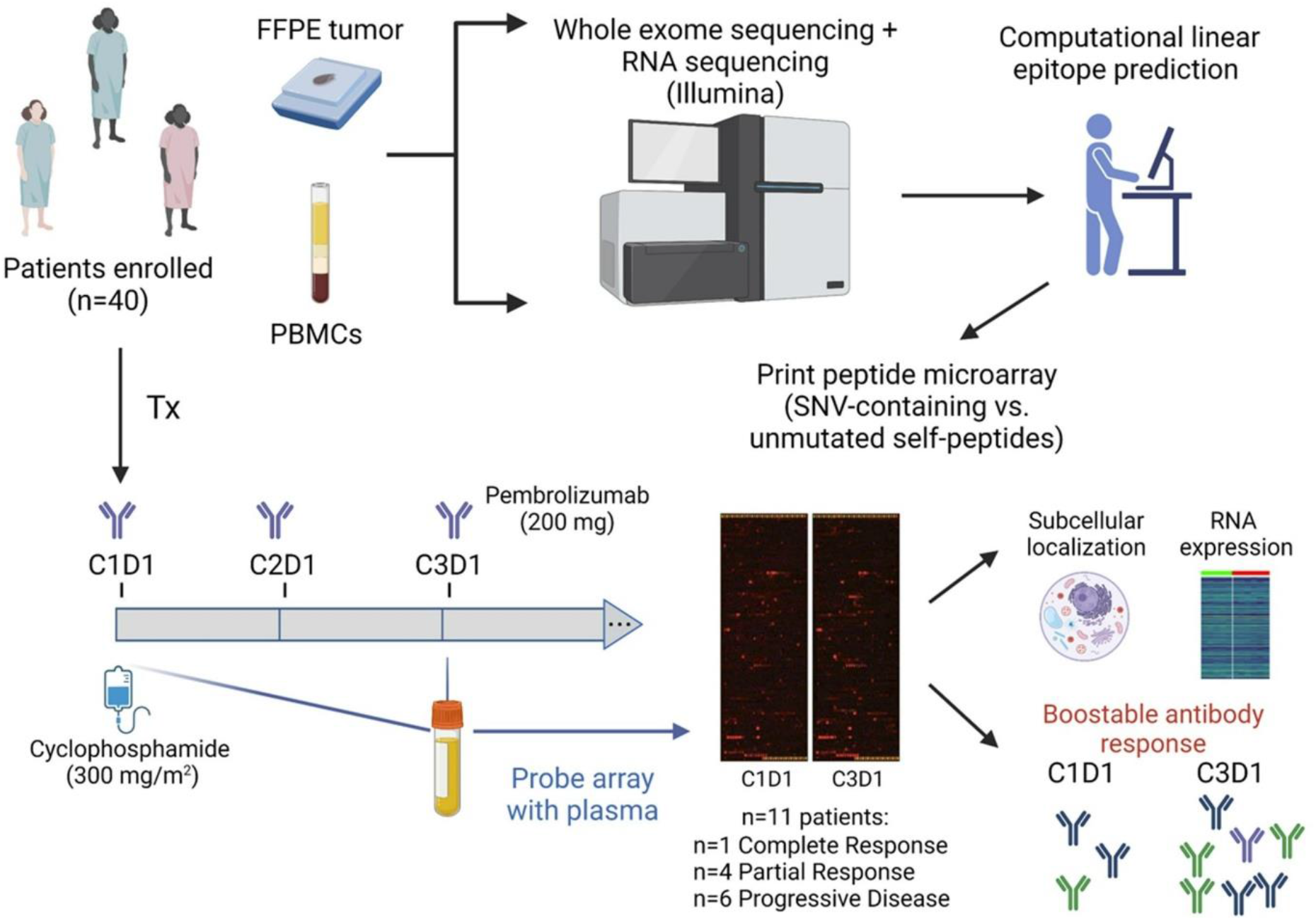
Experimental approach to examine neoantigen-specific antibody responses. Patients with metastatic triple negative breast cancer underwent a clinical trial to examine the efficacy of regulatory T cell depletion with cyclophosphamide plus PD-1 inhibition with pembrolizumab [10]. Eleven patients from this cohort were selected based on clinical response for analysis of tumor antigen-specific antibody responses via multiplex ELISA (peptide arrays). Downstream analyses included examination of protein subcellular localization and RNA expression, as well as antibody boostability relative to immunotherapy treatment. Antibody responses were seen at baseline, and in some patients, increased after therapy.

### Subcellular distribution and RNA expression analysis of peptides and associated antibody signal

We sought to understand whether there were any subcellular locations enriched for antibody signal-associated peptides. In order to focus on antibody signal with putative biological function, we compared antibody signal that was weak but above background noise (>500 F.U. and <2,000 F.U.) to antibody signal that was at least moderate in intensity (>2,000 F.U.). Subcellular localization analysis showed that peptides associated with antibody signal were most often derived from nuclear proteins, with cell surface and cytoplasmic subcellular locations also being more highly represented relative to membrane, mitochondrial and secreted proteins (**Fig. 2A**). Additionally, there was no subcellular distribution difference between peptides associated with weak versus stronger signal intensity (**Fig. 2A**). We were also interested in determining if parent proteins of peptides associated with stronger antibody signal intensity would have increased tumor RNA expression levels, but no such difference was observed (**Fig. 2B**).

**Figure 2.**
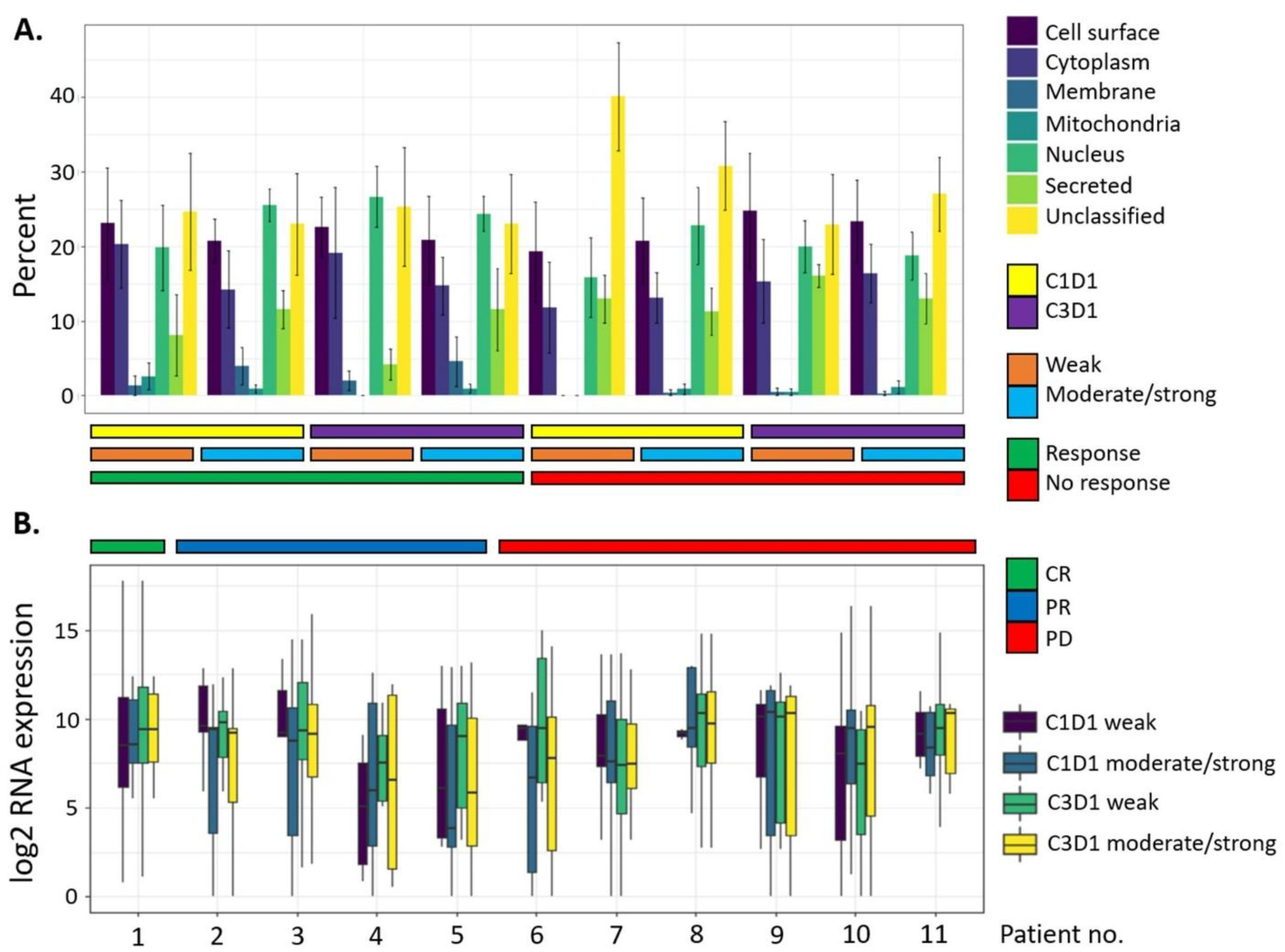
Subcellular protein localization and RNA expression do not associate with antibody signal or response group. **(A)** Subcellular localization of parent proteins of self/mutant peptide pairs subset based on associated antibody signal level (weak vs. moderate/strong), and further categorized by treatment time point and response. Peptide pairs were considered to have moderate/strong signal if either peptide of the pair (self or mutant) had >2,000 F.U. signal intensity. Peptide pairs were considered to have weak signal if both peptides of the pair (self and mutant) had signal intensity >500 F.U. and <2000 F.U. Error bars represent standard error (n=5 for responders; n=6 for non-responders). Subcellular localization was determined using the R Bioconductor package SubCellBarCode (https://bioconductor.org/packages/release/bioc/html/SubCellBarCode.html) and the R package UniprotR (https://rdrr.io/cran/UniprotR/). Each gene was assigned a subcellular annotation based on consensus between the outputs of these two R packages. If there was no consensus, then the subcellular annotation attained using SubCellBarCode was used as this method is based upon subcellular confirmation using mass spec data from 5 cell lines. For annotation of cell surface proteins, the Cancer Surfaceome Atlas (doi: 10.1038/s43018-021-00282-w) was used. **(B)** Distribution of RNA expression of parent proteins categorized according to antibody signal level, treatment time point and response.

### Observation of patient-specific patterns of antibody boostability

We examined the relationship between pembrolizumab treatment and boostability of antibody signal (e.g., increase in antibody signal between pre-treatment and post-treatment time points). ICB-dependent boost of both self-peptide-specific and mutated peptide-specific antibody signal was observed for some patients, but this was not significantly associated with clinical response class (**Fig. 3**). Patients 1 (CR) and 2 (PR), which were the patients with the highest tumor mutation burden, exhibited strong ICB-dependent antibody boost. The boost observed with patient 1 was largely driven by an increase in antibody signal to self-specific peptides (**Fig. 3B-D**), although mutated peptides were also boosted. Alternatively, the boost observed with patient 2 was driven by an increase in non-specific antibody signal (e.g., antibody response to both mutant and matched self-peptides). Of note, signal boost was not associated with clinical response to checkpoint inhibition, although the magnitude of boost was generally higher in responders than in non-responders. To gauge the absolute value of antibody signal boost relative to ICB treatment, we calculated the difference between total antibody signal at C3D1 and C1D1 (**Fig. 3C-D**). Interestingly, patient 1 exhibited the strongest absolute signal intensity difference relative to immunotherapy treatment. Together, these data suggest a possible association between ICB-dependent antibody boostability and clinical response.

**Figure 3.**
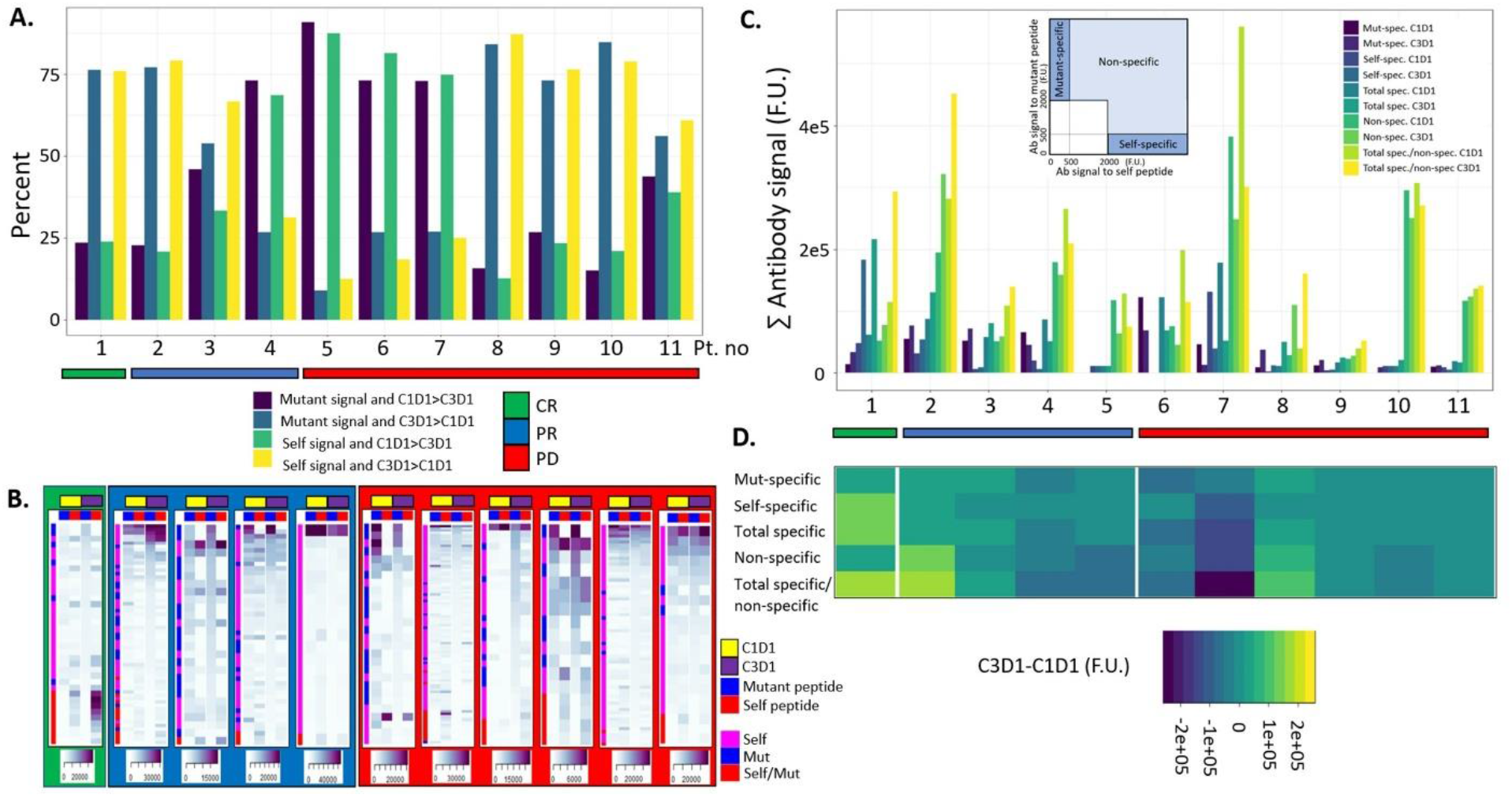
Boostability of antibody response relative to ICB treatment. **(A)** Percentage of mutant or self-peptides that had associated antibody signal (>500 F.U. at either C1D1 or C3D1) and greater signal relative to the other time point. **(B)** Heatmap depiction of antibody signal relative to treatment time point. Signal was ranked according to intensity of signal to C3D1 mutant peptide. A peptide pair was included if there was any signal >2,000 F.U. for either self or mutant peptide at either C1D1 or C3D1 time point. Colored sidebar denotes whether antibody signal was associated with mutant peptide, self peptide or both. **(C)** Absolute antibody signal (summed) categorized by signal specificity, treatment time point and response. Inlay depicts signal thresholds that were used to denote specificity classes. **(D)** ICB-associated boostability difference in antibody signal, which is derived by subtracting C1D1 antibody signal from C3D1 signal for respective specificity classes.

### Boostable self-specific antibodies to EPS8 dominate antibody response in complete responder

We next focused on the ICB-dependent boost that was observed in the complete responder to try and determine if there was something particular about this boost that associated with clinical response. A strong boost to EPS8 self-peptides was seen (**Fig. 4A**), and these peptides were located within an N-terminal region of the protein corresponding to residues S187-P107 (**Fig. 4B**). This region was predicted to have a mixed linear/helical structure, be surface-exposed/solvent-accessible, and to be disordered/flexible (**Fig. 4C-F**). These physical properties may contribute to a potentiated B cell response against these epitopes.

**Figure 4.**
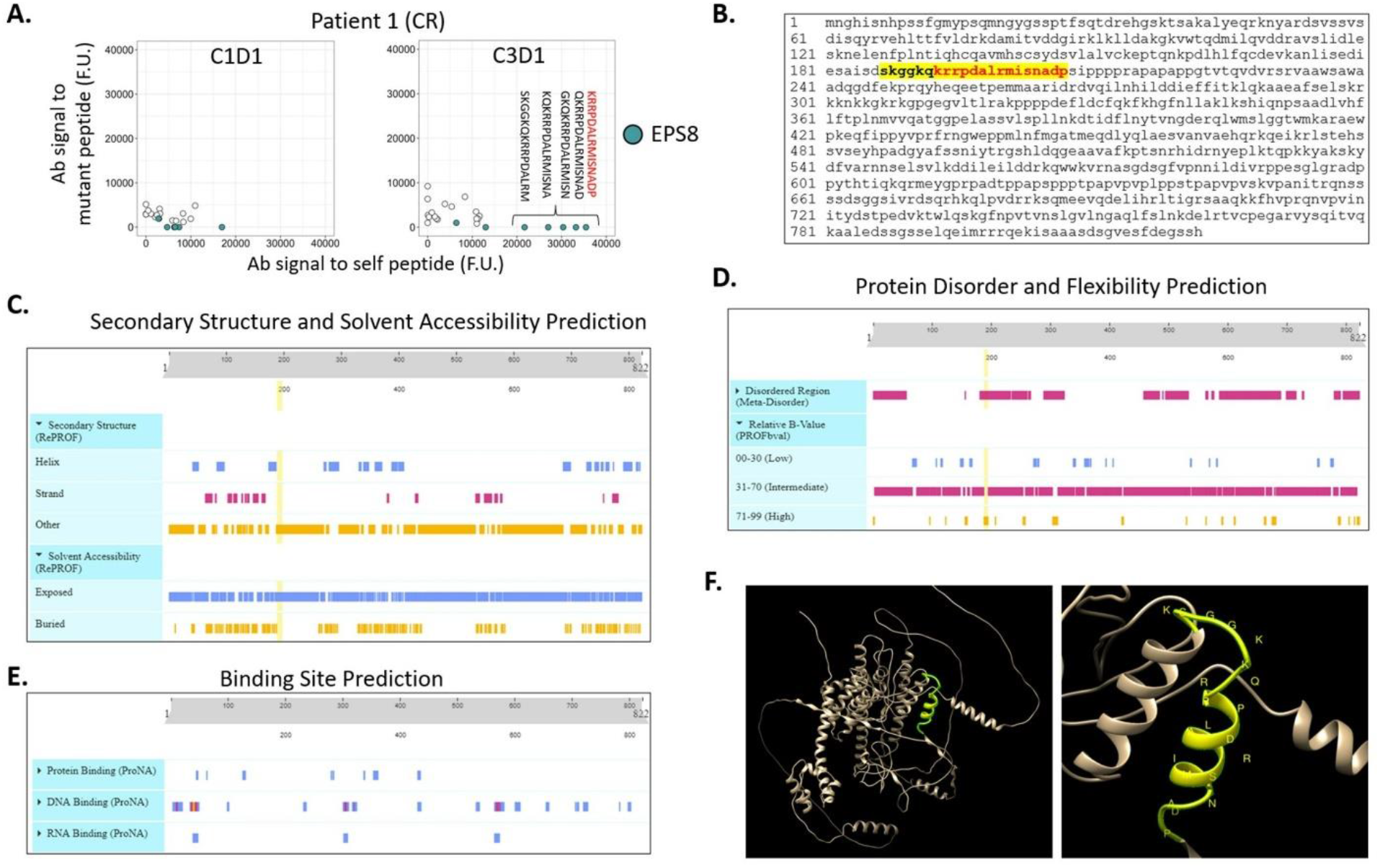
Complete responder displayed ICB-dependent boostability of both mutant and self-peptides, with particularly strong boost of IgG specific to native EPS8 peptides. **(A)** Relative antibody signal to self-versus mutant peptides at C1D1 and C3D1. Annotation shows ICB-dependent boost of antibody signal to self-EPS8 peptides. Peptides with greatest boost corresponding to S187-P107 residues are shown. **(B)** Primary protein structure of EPS8, with S187-P107 highlighted. The primary structure for EPS8 protein was input into PredictProtein (https://predictprotein.org) to query secondary structure and solvent accessibility **(C)**, protein disorder and flexibility **(D)**, and macromolecular binding site predictions **(E). (F)** Tertiary structure of EPS8 as predicted by Alphafold Protein Structure Database (https://alphafold.com), with S187-P107 highlighted in yellow. 3D structure was visualized and annotated using Chimera v1.16 software (https://www.cgl.ucsf.edu/chimera/).

## DISCUSSION

T cells dominate our concept of “The Cancer Immunity Cycle” [1] for good historical reasons: they kill tumor cells directly, have efficacy in adoptive transfer, and associate with clinical response to immune checkpoint inhibition. That said, studies from our group and others over the past 5 years have implicated the B cell arm of the adaptive immune system in tumor control. In breast cancer specifically, B cell population features consistent with antigen-driven clonal expansion associate with improved survival and response to immunotherapy. Thus, there is a need to discover the action(s) of tumor antigen-specific B cells in anti-tumor immunity in breast cancer.

We have taken a step in that direction by measuring tumor antigen-specific antibodies in the plasma of breast cancer patients treated with immunotherapy. There is evidence that anti-tumor antibodies can be important to achieve curative responses in large established murine tumors [24], and deeper understanding of the intricacies of antigen-specific B cell responses is necessary. Importantly, this study establishes that ICB can boost antibody responses to neoantigens, thus providing rationale for combining B cell-epitope-targeting vaccine strategies with ICB. Moreover, in the case of the complete responder, we observed a boostable antibody response targeting EPS8 protein, which is a known oncogene potentiating growth/survival (e.g., mTOR/PI3K/AKT/EGFR signaling) in multiple cancer types including breast cancer [25]. This finding highlights the potential utility of immunotherapeutic strategies aimed at boosting antibody responses to tumor-associated antigens.

The present study is limited in important ways. Due to the relative lack of responders in the study cohort (n=6 responders out of n=40 patients enrolled; 1 PR had data that was unusable due to low signal-to-noise ratio), this study is not powered to associate antibody signal with clinical response or genomics and immunogenomics features between patient groups. It also lacks power to interrogate the relative frequency of mutation-specific, unmutated self-specific and cross-reactive antibodies in the treated population. Metrics of IgG abundance/boostability were not corrected for TMB, and such analysis may yield further insight in later studies with larger cohorts. The framework of this analysis does not provide evidence that these antibody signals are associated with tumor growth, cytotoxicity, support of T cells, or other biological function(s). It is possible that the observed antibody boostability is merely a surrogate of T-cell expansion that may or may not have a therapeutic effect. Even if the observed antibody responses are real, we do not know how generalizable antibody production is across patients or tumor types. An additional deficit of this study is that it is limited in antibody discovery, as we have neither predicted nor measured conformational B cell epitopes, which may be the dominant epitopes in the system. It is unclear how these antibodies, or others targeting autoantigens, contribute to immunotherapy response.

In summary, we have performed an initial analysis of antibody responses in triple negative breast cancer patients treated with pembrolizumab following cyclophosphamide. We have found both tumor neoantigen-specific, self-specific and nonspecific antibodies that increased in signal after two cycles of pembrolizumab therapy. Similar studies with increased power to delineate the characteristics and contribution of ICB-dependent antibody responses to clinical response are warranted.

## LIST OF ABBREVIATIONS

DCs: Dendritic cells
EPS8: Epidermal Growth Factor Receptor Pathway Substrate 8
ICB: Immune checkpoint blockade
TLS: Tertiary lymphoid structure
TNBC: Triple negative breast cancer

## DECLARATIONS

### Contributors

EDR and BGV conceived the study. MGW performed in silica peptide predictions. ER performed all data analysis and figure preparation. EDR and BGV wrote the manuscript. All authors contributed to the review of the manuscript.

### Ethics approval

This study was approved by UNC IRB #16-1025.

### Data availability

Genomics data are available in a public open access repository.

## Acknowledgements

The authors thank Ken Fowler and Karen McKinnon with the Immune Monitoring and Genomics Facility (IMGF) at UNC-Chapel Hill for their assistance with this study. We also thank the patients in this study and their families, without whom this study would not have been possible.

